# The splicing factor kinase SRPK1 is a therapeutic target for Peripheral Vascular Disease

**DOI:** 10.1101/2024.04.17.589996

**Authors:** Sohni Ria Bhalla, Mussarat Wahid, Jason O Amartey, Amira Zawia, Federica Riu, Yizhuo Gao, Jyoti Agrawal, Amy P Lynch, Maria JC Machado, Tom Hawtrey, Ryosuke Kikuchi, Kathryn R Green, Lydia Teboul, Claire Allen, Zoe Blackley, Keerthana Rajaji, Daisy Marsden, Jennifer Batson, Steven J Harper, Sebastian Oltean, Winfried Amoaku, Andrew V Benest, Jonathan C Morris, Bruce Braithwaite, David O Bates

**Author notes:** Corresponding author: David Bates, Centre for Cancer Sciences, School of Medicine, Biodiscovery Institute, University of Nottingham, Science Road, NG7 2RD.

## Abstract

**Introduction:** In peripheral arterial disease (PAD) anti-angiogenic VEGF-A_165_b isoform overexpression in monocytes contributes to impaired collateralisation. Serine-arginine protein-kinase-1 (SRPK1) regulates VEGF splicing. To determine whether SRPK1 controlled monocytic VEGF, impairing collateralisation, we investigated SRPK1 inhibition and monocyte-specific knockout in mouse models of and in human monocytes from PAD.

**Methods:** VEGF-A_165_b activity was measured by co-culture of PAD patients’ monocytes with endothelial cells with SRPK1 inhibition. Mice with impaired revascularisation due to soluble-frizzled-related-protein-5 knockout (Sfrp5^-/-^), monocyte-specific Wnt5a gain-of-function (LysM-Wnt5a^GOF^), or obese mice on a high-fat high-sucrose (HF/HS) diet were subjected to femoral artery ligation and treated with SRPK1 inhibitor. We generated an SRPK1 conditional knockout and crossed it with monocyte-specific (LysM-Cre) driver line to specifically knockout SRPK1 in monocyte lineages. Blood flow was measured by Laser Speckle Imaging before, and for 28 days after surgery.

**Results:** Monocytes from PAD patients significantly inhibited endothelial cell migration, which was reversed by an anti-VEGF-A_165_b antibody. Surprisingly, migration was stimulated by SRPK1 inhibition, switching splicing from VEGF-A_165_b to VEGF-A_165_a. In Sfrp5^-/-^, LysM-Wnt5a^GOF^ and HF/HS mouse models of PAD, blood flow was improved by SRPK1 inhibition. Impaired revascularisation in LysM-Wnt5a^GOF^ mice was rescued in LysM-Wnt5a^GOF^:SRPK1^MoKO^ mice, which had a phenotypic shift towards M2 macrophages. Impaired blood flow recovery was also rescued in obese-SRPK1^MoKO^ mice.

**Conclusion:** VEGF splicing in monocytes is regulated differently from VEGF splicing in epithelial or cancer cells suggesting that control of splicing is dependent on cell type and/or environment. SRPK1 inhibition enhances collateralisation in mice, and in human *in vitro* models of monocyte-dependent impaired angiogenesis.

**New and Noteworthy:** A novel potential treatment for peripheral arterial disease (PAD) is described. Inhibition of SRPK1, or knockout in monocytes, induces angiogenesis by preventing splicing to anti-angiogenic VEGF (VEGF-A_165_b) in patients and animal models. In PAD, monocyte splicing control is different from other cell types and SRPK1 inhibition by drug like compounds can alter macrophage phenotype and reverse PAD in mice using a new cell specific SRPK1-LoxP mouse.

## Introduction

More than 4 million people in the UK have diabetes (7.9% of the population), predominantly type 2 (T2D). This is predicted to rise by 20% by 2030 (data from APHO Diabetes Prevalence Model). Progressive atherosclerosis and microvascular disease are the leading causes of amputations and are most common in people with diabetes. However, in otherwise healthy individuals, progressive atherosclerosis and microvascular disease can be circumvented by the development of collateral vessels. This collateral formation is impaired in people with diabetes, resulting in more severe PAD, critical limb threatening ischemia (CLTI) and ischemia of other tissues, including coronary vascular disease (CVD), leading to increased incidence of fatal myocardial infarction (MI)(1). Understanding why diabetes impairs collateral formation, and identifying ways that can overcome that inhibition, could provide radical, effective new therapies for people with diabetes, PAD and CVD.

Hyperactivation of the Wnt5a pathway in monocytes has been proposed to be a key mechanism through which metabolic syndrome patients(2) have impaired collateralisation(3–5). In T2D and in obesity circulating monocytes are pro-inflammatory(6). These circulating monocytes have increased VEGF production(7), suggesting that they should be able to aid arteriogenesis and angiogenesis and stimulate collateral formation. However, the monocytes appear unable to elicit the actions of VEGF(8). This paradox – increased VEGF production but reduced angiogenic capability in diabetic patients – may be partly explained by findings that monocytes from T2D patients, and mouse and rat models of T2D, have increased Wnt5a expression and activity(4). Sfrp5, the endogenous inhibitor of Wnt5a is reduced in plasma from patients with obesity, T2D or both(9, 10). Sfrp5 knockout (SFRP5^-/-^) in adult mice significantly impaired recovery of blood flow and angiogenesis after hind limb ischemia (HLI), a model of collateralisation in mice. This impairment is reproduced by a Wnt5a overexpressing monocytic lineage-specific gain of function animal (LysM-Wnt5a^GOF^). VEGF-A mRNA and protein levels are dramatically upregulated in both these mouse strains after ischemia, contrary to the findings of reduced angiogenesis(5).

VEGF-A exists as two functionally contrasting families of isoforms resulting from mRNA processing(11). Alternative 3’ splice site selection in the terminal exon, exon 8, can generate either pro-angiogenic isoforms termed VEGF-A_xxx_a where xxx is the number of amino acids in the mature polypeptide (e.g. VEGF-A_165_b), or isoforms that can inhibit angiogenesis by use of a splice site 66 nucleotides downstream of the proximal 3’ splice site, termed VEGF-A_xxx_b(12). VEGF-A_165_b is the most studied member of this family. VEGF-A_165_b acts as a partial agonist for its receptors(13–15), inhibiting VEGF-A_165_a induced angiogenesis(15), but protecting endothelial and epithelial cells (including ocular and renal epithelial cells) from cytotoxic insults(16, 17). VEGF-A_165_b binds to the tyrosine kinase receptors VEGFR1 and VEGFR2(13–15) but not neuropilin or heparan sulphate proteoglycans(14) resulting in partial activation of VEGFR2(18), unstable kinase activation due to lack of phosphorylation of Tyr1054 in VEGFR2(13) and inhibition of VEGFR1 signalling(19).

Interestingly, the impaired revascularisation seen in Sfrp5^-/-^ and LysM-Wnt5a^GOF^ mice is VEGF-A_165_b dependent as VEGF-A_165_b neutralising antibodies reversed the blocked revascularisation(5). Moreover, VEGF-A_165_b is upregulated in both these strains after ischemia and in mouse obesity models (ob/ob mice and wild type C57Bl6 mice fed a high fat, high sucrose diet for 12 weeks), and VEGF-A_165_b selective antibodies also improved collateralisation in both obese models with peripheral ischemia(5, 19–21) and in normal mice with coronary vascular disease(22). VEGF_165_b has also been shown to be upregulated in people with peripheral arterial disease (PAD) and in people with coronary vascular disease (CVD)(23), both at the RNA level in circulating monocytes(5), and at the protein level in plasma(22), muscle and macrophages(5). These results provide a potential explanation for impaired collateral formation in diabetes, and a potential novel therapeutic strategy to reduce cardiovascular disease in diabetic patients. That strategy would require selective inhibition of VEGF-A_165_b in monocytes in people with diabetes and ischemia.

Pre-mRNA splicing is orchestrated by splicing factors. In contrast with constitutive splicing, alternative splicing refers to the recruitment of a unique pattern of splicing factors (tissue and developmental stage-specific) resulting in transcripts that encode different proteins, often with contrasting properties. The actions of key splicing factors (SFs) such as the SR proteins SRSF1 and SRSF6 can be modulated by small molecular weight inhibitors of their cognate kinases SRPK1/2 (e.g. SPHINX31(24)) and CLKs (e.g. TG003)(25). SRSFs are known to affect VEGF-A splicing – in epithelial cells, SRSF6 over-expression switches expression to VEGF-A_165_b(26), and SRSF1 to VEGF-A_165_a in kidney and ocular epithelial cells(27, 28), and SRSF2 to VEGF-A_165_a in lung cancer cells(29). In epithelial and cancer cells, inhibition of SRSF1 phosphorylation by SRPK1 blockade inhibits VEGF-A proximal splice site selection in exon 8(30, 31), resulting in less VEGF-A_165_a, and SRPK1 inhibition is an effective anti-angiogenic strategy for retinal neovascularisation and choroidal neovascularisation in eye disease(30, 32, 33). However, we do not yet know what regulates VEGF splicing in monocytes, or whether this holds true in monocytes from diabetic patients with cardiovascular disease, or monocytes as they differentiate into macrophages during adherence and extravasation.

We therefore investigated the mechanism of splicing control in monocytes, and show that SRPK1 in monocytes, in contrast to epithelial cells, switches splicing to the VEGF-A_165_b isoform in monocytes, and SRPK1 inhibition is able to reverse peripheral vascular disease in mice, and induce the angiogenic capability of human monocytes.

## Methods

### Patient samples

Patients with peripheral artery disease (ankle brachial pressure index <0.9) were enrolled from Queen’s Medical Centre, Nottingham (UK) after obtaining their written informed consent under ethics number IRAS265512 between October 2019 and June 2022. As this was not a clinical trial, patient data was fully anonymised with only entry criteria (ABI<0.8, age >50) recorded, and samples destroyed after use. All studies were performed in accordance with the Declaration of Helsinki. 50 mL of whole peripheral blood was collected in EDTA tubes. Peripheral blood mononuclear cells (PBMCs) were isolated from blood by density gradient centrifugation using Ficoll® Paque Plus (Cytiva, Marlborough, MA, USA). Blood was buffered by mixing it with an equal volume of PBS. 15mL of Ficoll-Paque Plus solution (Cytiva) in a 50 mL conical tube was carefully overlaid with 30 mL of diluted blood and centrifuged for 20 minutes at 1,000 g at room temperature with slow accelerator and no brake. The interface containing the cellular layer was collected from each tube with a transfer pipette, washed with PBS and placed into a 50 mL tube and centrifuged for 10 minutes at 600g with maximum acceleration and deceleration at room temperature. The obtained PBMCs were treated with 5mL of red blood cell lysis buffer (0.83g NH4Cl, 0.1g KHCO3, 5% EDTA dissolved in 100 mL of distilled water and filter sterilized using 0.22 µM filter) and incubated for 5 minutes at room temperature to lyse the remaining red blood cells. This tube containing lysed cells was then centrifuged for 6 minutes at 400g at room temperature. The PBS wash was repeated followed by centrifugation for 6 minutes at 400 g at room temperature, cell count and viability assessment were performed. To separate monocytes from the PBMCs, antihuman CD14 MicroBeads (Miltenyi Biotec, Bergisch Gladbach, Germany), were used as per manufacturer’s protocol. The CD14 positive cells were then re-suspended in RPMI 1640 medium (Fisher Scientific, Leicester, UK) supplemented with 10% human AB serum (Zen Bio, Durham, NC, USA).

### Endothelial cell migration assay

Human Umbilical Vein Endothelial Cell (HUVEC) (PromoCell GmbH, Heidelberg Germany) were serum starved overnight in Endothelial cell basal medium (ECBM) supplemented with endothelial cell growth kit (PromoCell GmbH, Heidelberg Germany) without serum. Transwell migration assays were performed in 24-well Thincert, (8.0µm pore size; Greiner Bio-One, GL, UK) placed in 24-well plates containing 0.1% fetal calf serum without or with 40ng/ml VEGF_165_a, 40ng/ml VEGF_165_a and 40ng/ml VEGF_165_b, or both isoforms and increasing concentrations of mouse anti-VEGF-A_165_b antibody(15), or in human AB serum (HSER-ABP100ML AMSBio, Oxford, UK) in RPMI 1640 media (Fisher Scientific, Leicester, UK) alone (0.5% control), with 40ng/ml VEGF_165_a or with 40ng/ml VEGF_165_a CD14^+^ labelled monocytes suspended at a density of 5×10^4^ cells and treatments as shown. HUVEC were suspended in serum-free ECBM (without supplements) and plated at a density of 5×10^4^ cells on the upper chamber of the 8µm inserts. Endothelial cells were allowed to migrate overnight at 37°C under 5.0% CO_2_. The inserts were washed 3 times in PBS and fixed with 4% paraformaldehyde (PFA) for 15 minutes. After fixation, inserts were washed 3 times in PBS and stained with DAPI (nuclei stain) for 15 minutes at room temperature. Stained membranes were removed from inserts and mounted on the glass slide. The migrated cells were counted under a microscope in 3 different views away from the insert edge by an observer blinded to treatment (10x magnification). Each monocyte sample was reproduced in triplicate.

### Animal model

Sfrp5^-/-^ and LysM-Wnt5a^GOF^ mice were a kind gift of K Walsh. Mice with SRPK1 knockout were generated in collaboration at MRC Harwell by flanking exon 7 of *Srpk1* gene with LoxP sites. The promoter driven L1L2-Bact-P cassette was removed by crossing mice with flipase mediated recombination generating SRPK1^fl/fl^ mice. To generate the myeloid derived cell specific SRPK1 knockout mouse model (hereafter referred to as SRPK1^MoKO^), the Srpk1^fl/fl^ mice were crossed with Lysozyme M-Cre mice. To generate the double transgenic mouse model (hereafter referred to as LysM-Wnt5a^GOF^: SRPK1^MoKO^), the LysM-Wnt5a^GOF^ mice were crossed with the SRPK1^fl/fl^ mice. All mice were maintained on a C57/BLJ background. Mice were fed either a normal chow diet (Teklad global, 2018S) or a HF/HS diet (Bio-Serv, S1850) as indicated. The composition of the HF/HS diet was 35.8% fat, 36.8% carbohydrates and 20.3% protein. For the HF/HS diet, mice were maintained on a HF/HS diet from the age of 4 weeks old for the duration of the study (15-16 weeks).

### Hind limb ischemia surgical procedure

All animal experiments were conducted in accordance with the Animal Scientific Procedures Act (ASPA) of 1986 under a UK Home Office License at the University of Nottingham Biological Services Unit. All studies performed conform with the guidelines from the Directive 2010/63/EU of the European Parliament and the NIH Guide for the Care and Use of Laboratory Animals. Male and female (50:50) transgenic C57/BLJ mice were subjected to unilateral hindlimb ischemia between 10 to 12 weeks of age. Anaesthesia was induced with 2% isoflurane in 100% oxygen at rate of 2L/min. Body temperature was controlled throughout using a homeothermic blanket and rectal probe (Harvard Apparatus). The hair was removed from the hind limb using Nair hair removal cream and sterilised using chlorohexidine-based solution. Mice were administered a pre-operative analgesic, buprenorphine (0.05mg/kg, subcutaneous) and saline solution (0.9% sodium chloride at 40ml/kg). An incision was cut in the left medial thigh and the connective tissue was teased apart to expose the femoral artery, vein and nerve. The nerve bundle and vein were teased apart from the femoral artery. The femoral artery was ligated above and below the epigastric branch and electro-coagulated in between to induce ischemia to the left hind paw. The blood flow to both paws was measured using the laser speckle imaging system (Moors FLIP2, Moors instruments). The blood flow was monitored on pre-operative day 0 and post-operative days 0, 3, 7, 14, 21 and/or 28. The ratio of the blood flow was calculated between the ischemic vs contralateral paw to measure the blood flow recovery throughout the duration of the study to account for day to day variation in blood flow due to potential changes in body temperature, ambient temperature, central blood pressure, etc. If the reduction in blood flow was less than 70% the animals were excluded for subsequent analysis *a priori*. For SPHINX31 *in vivo* treated experiments, animals were treated with 0.8mg/kg SPHINX31 synthesised as described in (24) bi-weekly for the duration of the study.

Twenty Sfrp5^-/-^ and twenty wild type mice were used, and 8 of each type treated with SPHINX31. Two sfrp5^-/-^ and one wild type animal were excluded due to lack of impaired flow (as outlined above). Eight of each group were perfuse fixed for immunofluorescence and four had protein extracted for VEGF assessment. In one wild type animal protein was lost during processing.

Eighteen Wnt5a^GOF^ mice were used, 9 in the control group and 9 in the SPHINX31 treated group. 2 mice were excluded from the Wnt5a^GOF^ and one from the SPHINX31 group due to insufficient ligation.

### Immunofluorescent staining

At the experimental end-point, mice were euthanized by cardiac perfusion with PBS, followed by 4% PFA/PBS and the gastrocnemius muscle was collected. Tissue was post fixed in 4% PFA/PBS overnight at 4°C, cryoprotected in 30% sucrose and embedded in OCT. The muscles were cryosectioned into 5μm-thick sections and stained for IB_4_ (1:150, Sigma), αSMA-Cy3 (1:300, Sigma), M1 marker iNOS (rabbit PA1-036), rabbit anti-SRPK1 (140731-AP, 1:150) both ThermoFisher, M2 marker CD206 (abcam ab 64693, 1:200) or humanised anti-VEGF_165_b (1:50, Wahid et al submitted). For immunofluorescent staining, a standard protocol was followed. In brief, sections were fixed in ice cold acetone for 5 minutes before permeabilising with 0.5% Triton X-100 for 20 minutes. Sections were then blocked (4% BSA/PBS with 0.5% Triton-X100) for 1 hour followed by incubation with primary antibodies overnight. Appropriate Alexafluor secondary antibodies were used in 1:200 dilution for 1 hour then the sections were mounted with mounting media containing DAPI. Sections were imaged using a confocal microscope and fluorescent microscope. Double positive macrophages (F4/80^+^iNOS^+,^ F4/80^+^CD206^+^ and F4/80^+^ SRPK1^+^) were counted in 5 field of views at X100 magnification using double thresholding and overlay method in FIJI.

### Isolation of CD11b mouse monocytes

Freshly isolated bone marrow and spleen from 10 to 12 weeks of age WT and SRPK1^MoKO^ mice were used. Bone marrow immune cells and splenocytes were exposed to red blood cell lysis. The remaining cells were separated from a mixed population of cells to monocytes and non-monocytes using the CD11b monocyte isolation kit (Miltenyi Biotech) following the manufacturer’s instructions using Miltenyi LS columns.

### Western blotting

Mouse gastrocnemius muscle tissue samples obtained on postoperative day 7 were homogenized in lysis buffer containing 20 mm Tris-HCl (pH 8.0), 1% Nonidet P-40, 150 mm NaCl, 0.5% deoxycholic acid, 1 mm sodium orthovanadate, and protease & phosphatase inhibitor cocktail (Thermo Scientific). Protein content was determined by the Bradford method. Equal amounts of protein lysates were resolved by SDS-PAGE. The membranes were immunoblotted with the indicated antibodies (anti-SRPK1 antibody (ab38017), Abcam, anti-SRSF6 antibody (ab244425), and anti-SRPK1, HPA016431, Sigma Aldrich) at a 1:1000 dilution followed by the secondary antibody conjugated to horseradish peroxidase at a 1:1000 - 5000 dilution. An ECL western blotting Detection kit (Thermo Scientific) was used for detection. Immunoblots were normalized to total loaded protein.

### Cell Lysis

Protein was extracted from monocytes after treatment. Cells previously treated with inhibitors were centrifuged at 10,000 RPM at 4°C until a pellet was formed. Cell lysis buffer (1xNP40, 1mM phenylmethylsulfonyl fluoride (PMSF), 10mM Sodium orthovanadate (Na_3_VO4), 1x Protease Inhibitor cocktail (Roche), 10mM sodium fluoride) was added to the pellet for 10 mins. The cell extract was vortexed for 15 sec three times over a 10 min incubation on ice. Samples were centrifuged at 12,000 x g for 10 min at 4°C and the supernatant was collected in a fresh Eppendorf and stored at −80°C. Protein concentration was quantified using a Pierce BCA Protein Assay Kit (Thermo Scientific 23225).

### ELISA

High binding 96-well plates were coated with 100µl of either 0.75µg/ml hVEGF-A_165_a (ab14994) or 10µg/ml VEGF-A_165_b capture antibody overnight at room temperature on the shaker. The plates were washed three times with wash buffer (0.05% Tween-20/PBS, T-PBS) and blocked twice with 300µl/well of Superblock and discarded immediately. Serial dilutions were prepared with rhVEGF-A_165_a and rhVEGF-A_165_b diluted in PBS (ranging from 1.95 to 4000pg/ml as well as a blank and a negative control) were added in duplicate (100µl/well). 100µg of protein tissue was added in duplicate to each well. The 96-well plates were incubated for two hours at room temperature on a shaker. Then the 96-well plates were washed three times and incubated with 100µl of detection antibody from kit diluted in 1% BSA/PBS for two hours. After another wash step, 100µl/well of horseradish peroxidase (HRP)-conjugated streptavidin (diluted according to kit in 1% BSA in PBS was added and the plate was left in the dark for 30 minutes. After a final wash step, 100µl/well of the substrate solution (Abcam) was incubated for 20 to 30mins at room temperature in the dark. The reaction was stopped by the addition of 50µl/well of either 1M hydrochloric acid or sulphuric acid. The colorimetric reaction was measured using the microplate reader at 450nm with a 620nm reference (Infinite F50, Tecan UK). The protein concentration in each sample was calculated by subtracting absorbance values from the blank and the standard curves was used to calculate protein content in each sample.

### PCR

RNA was isolated from CD11b^+^ and CD11b^-^ cells using Trizol. 1μg RNA was transcribed to cDNA using the Takara Primescript RT kit (Takara) as directed by the manufacturer. Each 15μl consisted of 0.4uM primers, 2x GoTaq master mix (Promega) and 1μl cDNA. The primer sequences for the target genes were: *Srpk1*: forward, 5’-TCTCGCCATGGAGCGGAAAG-3’, reverse, 5’ TCCAGGCCCTGTAGCACTTG-3’; *Gapdh*: forward, 5’-CCATCTTCCAGGAGCGAGAC-3’, reverse, 5’-GCCC TTCCACAATGCCAAAG-3’. The thermal cycling conditions compromised of an initial denaturation at 95°C for 3 minutes, followed by 35 cycles at 95°C for 30s, annealing at 62°C for *Srpk1* or 58°C for *Gapdh* for 30s, 72°C for 1 min. This was followed by a full extension at 72°C for 5 mins. RT-PCR was carried out for VEGF-A_165_b and VEGF-A_165_a using isoform specific primers Forward:5’-GGCAGCTTGAGTTAAACGAAC-3’, Reverse: 5’-ATGGATCCGTATCAGTCTTTCCTGG-3’ as previously described (34) PCR products were run on 2-3% agarose gels with 25ng/ml ethidium bromide. The agarose gel was visualised on a UV transilluminator GelDocEZ system.

### Statistical analysis

Statistical analyses were completed on GraphPad Prism, Microsoft Excel, FIJI and Laser Speckle imaging analysis software (Moor’s Instruments). All data is presented as the mean ± SEM as indicated in the figure legends. In all cases p<0.05 was considered statistically significant, but where 0.06>p>0.05 exact p values are shown. Statistical tests were used according to normality tests in Prism, and equivalence of standard deviations, as described in the Prism Manual. Statistical tests are described where used.

## Results

To determine the effect of SRPK1 inhibition on monocyte VEGF-A_165_b we treated monocytes from patients with PAD with 3µM SPHINX31 (a selective SRPK1 inhibitor) overnight and measured VEGF-A_165_a and VEGF-A_165_b using isoform specific ELISAs(35),(15). VEGF-A_165_a expression was not detected in PAD monocytes, but surprisingly was significantly increased by SRPK1 inhibition (Figure 1A). In contrast, VEGF-A_165_b expression was significantly reduced by SRPK1 inhibition (Figure 1B). Monocytes from patients with dominance of VEGF-A_165_b RNA as measured by RT-PCR, were subjected to SPHINX31 treatment, which significantly increased the ratio of VEGF-A_165_a:VEGF-A_165_b RNA as measured by RT-PCR (Figure 1C). To determine whether this change in expression would functionally affect the ability of monocytes to induce angiogenesis we developed an *in vitro* assay for VEGF-A_165_b mediated inhibition of endothelial cell migration. Figure 1D shows that in the presence of 1nM VEGF-A_165_a, endothelial cell migration across a transwell was significantly increased compared with control, but this was significantly decreased by exposure to monocytes from patients with PAD. This inhibition was dose dependently reversed with a neutralising antibody to VEGF-A_165_b (Figure 1D). Figure 1E shows that treatment with an SRPK1 inhibitor blocks the inhibition of migration by PAD monocytes. These results indicate that PAD monocytes produce VEGF-A_165_b, sufficiently to inhibit endothelial cell migration towards 1nM VEGF-A_165_a and that SRPK1 treatment switches splicing and sufficiently to stop the inhibition of migration by VEGF-A_165_b under these circumstances. To determine whether this would be effective in vivo we used an animal model of PAD previously shown to be VEGF-A_165_b dependent(5).

**Figure 1.**
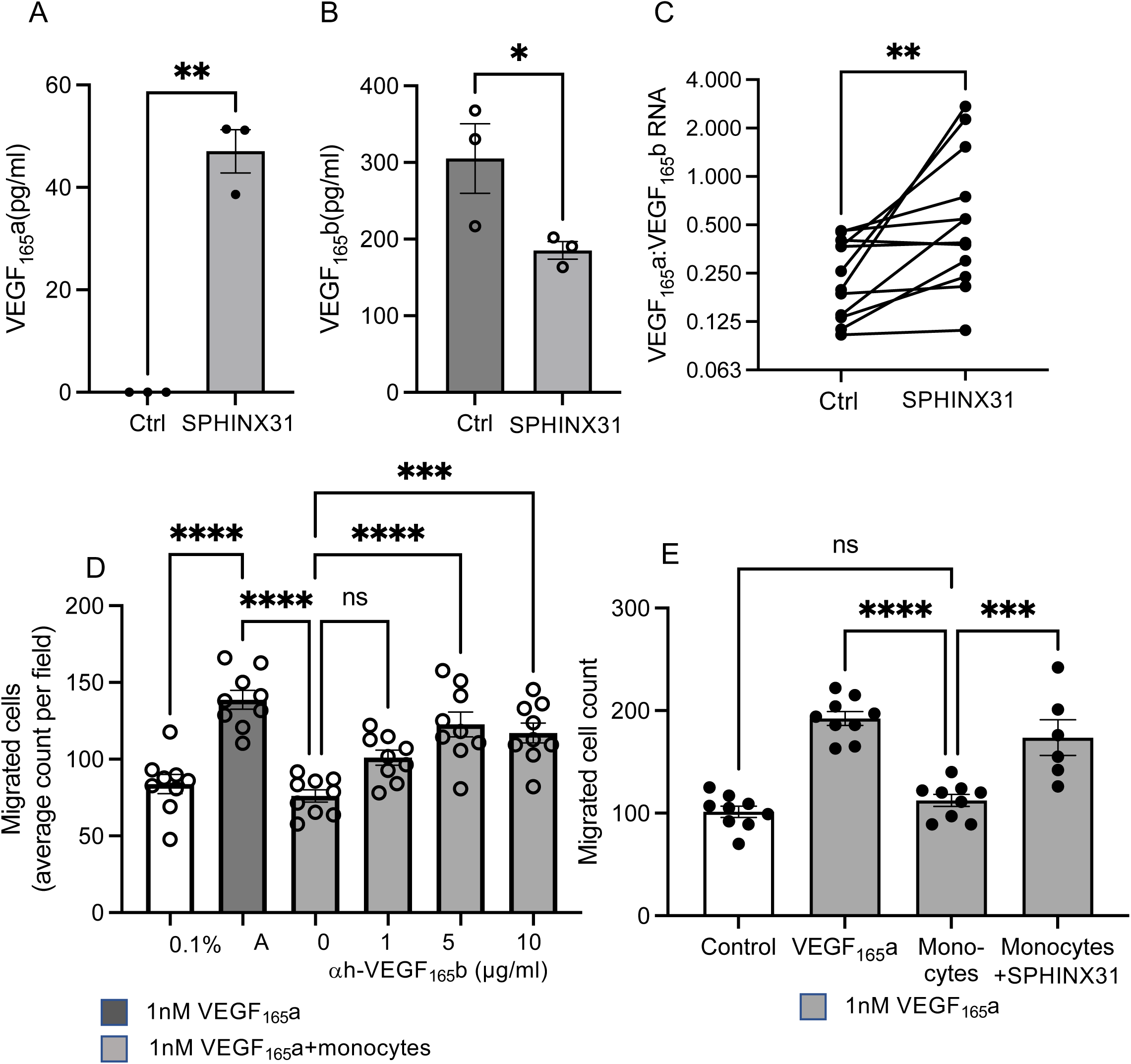
SRPK1 inhibition switches splicing to the angiogenic forms of VEGF in PAD monocytes. **(A)(B).** Monocytes from three patients with PAD were cultured for 24 hours in the presence of 3µM of SRPK1 inhibitor SPHINX31 or vehicle (0.1% DMSO). Protein was extracted and **(A)** VEGF-A_165_a or **(B)** VEGF-A_165_b measured by isoform specific ELISA. Data was analysed using a paired t-test (N=3 patients/group). **(C)** RNA was extracted from monocytes from 13 patients incubated with either vehicle or SPHINX31 and VEGF-A_165_b and VEGF-A_165_a measured by RT-PCR. Data was analysed using a paired t-test (N=12 patients/group). **(D)** Endothelial cells seeded onto the upper side of a transwell membrane and cultured with either media (0.1% serum control), 1nM VEGF-A_165_a, or 1nM VEGF-A_165_a and monocytes from nine PAD patients with increasing concentration of anti-VEGF-A_165_b antibody in the lower half of the transwell. After 12 hours the number of endothelial cells that had migrated were calculated. Data was analysed using a one-way ANOVA (N=9 patients/group). **(E)**. Endothelial cell migration in response to monocytes treated with 3µM SPHINX31. Data was analysed using a one-way ANOVA (three subjects, N=3 or 2 per subject). All results are shown as the mean ± SEM. *p<0.05, **p<0.01, ****p<0.0001.

### Impaired collateral revascularisation in Sfrp5 knockout (Sfrp5^-/-^) mice is reversed with SPHINX31, an SRPK1 inhibitor

Previous studies have identified that Sfrp5 acts to inhibit the non-canonical Wnt5a signalling pathway in macrophages and impaired revascularisation in a hindlimb ischemia model. This correlated with an upregulation of Wnt5a at a transcriptional level in the ischemic limb(4, 5), and upregulation of VEGF-A_165_b in monocytes(5). We therefore used this model to determine whether SRPK1 was able to modulate the angiogenic activity of monocytes in ischemia. Twenty Sfrp5^-/-^ and twenty wild type mice underwent left femoral artery ligation and the blood flow to the hind paws were monitored non-invasively using laser speckle imaging on pre- and post-operative days (Figure 2A). Quantitative analysis revealed that Sfrp5^-/-^ had impaired revascularisation compared to WT mice on days 21 (0.60 ± 0.04 vs 0.76 ± 0.04, p=0.007) and 28 (0.62 ± 0.04 vs 0.85 ± 0.08, p=0.0002) (Figure 2B). At the end of the study, a VEGF-A_165_b ELISA was performed on the gastrocnemius muscle which revealed that the Sfrp5^-/-^ mice had significantly greater VEGF-A_165_b protein content compared to WT mice in both ischemic and contralateral muscle (Figure 2C). The gastrocnemius muscle was collected, sectioned, stained and imaged for arterioles and capillaries (Figure 2D). The capillary density was unchanged in the ischemic gastrocnemius muscle of both mouse groups (650.1 ± 43.7/mm^2^ vs 721.9 ± 72.6/mm^2^, p=0.41) (Figure 2E). However, the arteriolar density was significantly reduced in the ischemic gastrocnemius muscle of Sfrp5^-/-^ mice (4.40 ± 0.90/mm^2^) compared to WT (7.01 ± 0.67/mm^2^, p=0.04) (Figure 2F). To determine whether SRPK1 and SR protein expression was altered in these mice, protein was extracted from total muscle tissue and subjected to immunoblotting for SRSF1, SRSF6 and SRPK1. SRSF1 was increased in both cytoplasm and nucleus of cells from muscle from Sfrp5^-/-^ mice, whereas SRSF6 was reduced. SRPK1 levels were increased in ischemia only in Sfrp5^-/-^ mice (Appendix Figure 1). To explore the significance of SRPK1 on revascularisation, SPHINX31(24),(36) was administered to Sfrp5^-/-^ mice via an intraperitoneal route after the ischemic hind limb surgery and bi-weekly thereafter. Quantitative analysis of the speckle images (Figure 2G) revealed that the flow was significantly increased in Sfrp5^-/-^ treated with SPHINX31 on post-operative days 14 (0.55 ± 0.04 vs 0.93 ± 0.08, p=0.001) and days 21 (0.51 ± 0.05 vs 0.75 ± 0.07, p=0.04) when compared to the Sfrp5^-/-^ treated with DMSO alone (Figure 2H).

**Figure 2.**
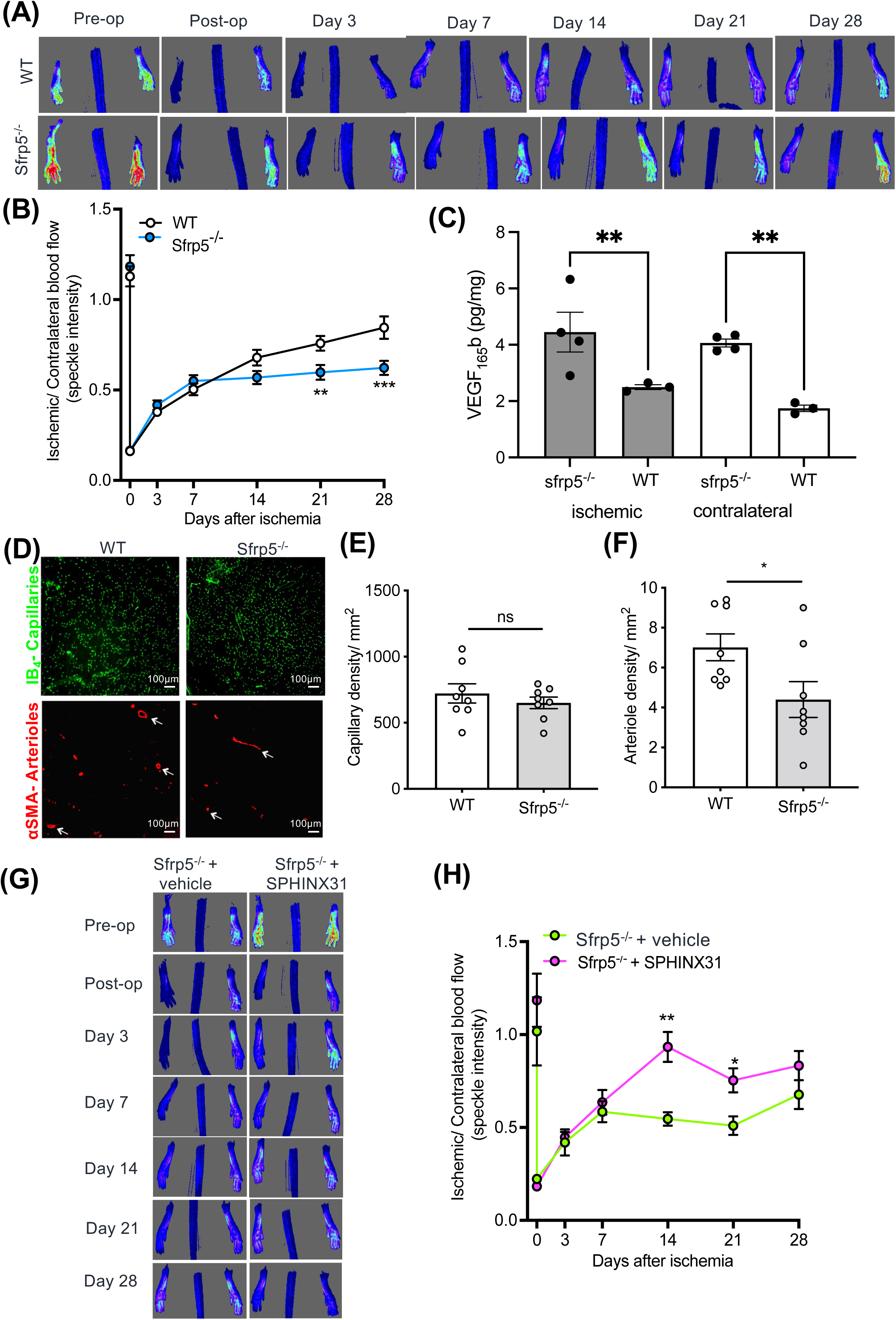
SRPK1 inhibition with SPHINX31 reverses impaired revascularisation in Sfrp5^-/-^ mice. **(A)** WT and Sfrp5^-/-^ mice underwent left femoral artery ligation and the blood flow to the paw was quantified using the laser speckle imaging system, as shown in the representative speckle images. **(B)** Quantitative analysis of the ischemic/ non-ischemic speckle intensity was calculated through to post-operative day 28. Data was analysed using a two-way ANOVA (N=19/group). **(C)** Measurement of VEGF-A_165_b from muscle tissue from Sfrp5^-/-^ and wild type mice. Sfrp5^-/-^ mice (N=4) had significantly greater VEGF-A_165_b expression than the wild type controls (N=3) in both ischemic and contralateral muscle. Data was analysed using a two-way ANOVA with Holm Sidaks (N=3/4/group). **(D)** Confocal images from the representative muscle sections of the gastrocnemius on post-operative day 28 stained with isolectin B4 for endothelial (green) and α-smooth muscle actin for vascular smooth muscle cells (red). Scale bar = 100 μm. **(E)** Capillary density was not changed in WT and Sfrp5^-/-^ mice. Data was analysed using an unpaired t test (N=8/group). **(F)** Arteriole density is reduced in Sfrp5^-/-^ mice when compared to WT mice. Data was analysing using an unpaired t-test (N=8/group). **(G)** Sfrp5^-/-^ mice received an intraperitoneal dose of DMSO or SPHINX31 (0.8mg/kg) bi-weekly for the duration of the study. The representative speckle images are shown on pre-operative day and on post-operative days 3, 7, 14, 21 and 28. **(H)** Quantitative analysis of the ischemic/ non-ischemic speckle intensity was calculated through to post-operative day 28. Data was analysed using a two-way ANOVA (N=7-8/group). All results are shown as the mean ± SEM. *p<0.05, **p<0.01, ***p<0.001.

### Inhibition of SRPK1 improves revascularisation in LysM-Wnt5a^GOF^ mice

To determine whether SRPK1 inhibition would exert the same effect in a different mouse model of Wnt5a activation, we used a monocytic Wnt5a overexpression (LysM-Wnt5a^GOF^) model. This resulted in a significant decrease in blood flow (Figure 3A) compared with wild type animals, which was reversed by SPHINX31 treatment. Quantitative analysis of the speckle intensity revealed that the blood flow recovery was significantly slower in LysM-Wnt5a^GOF^ mice when compared to WT mice on post-operative day 3 (0.19 ± 0.03 vs 0.72 ± 0.09 compared with the contralateral limb, p<0.0001), 7 (0.40 ± 0.03 vs 0.82 ± 0.10, p<0.0002) and 14 (0.59 ± 0.08 vs 0.92 ± 0.05, p=0.0039) but this was completely reversed by treatment with SPHINX31 as above (Figure 3B). Recovery between mouse groups was not different at post-operative day 21, when gastrocnemius muscle was collected, stained and imaged for capillary and arterioles (Figure 3C). The capillary density was significantly decreased in the ischemic gastrocnemius muscle of LysM-Wnt5a^GOF^ (311 ± 18mm^2^) when compared with WT mice (491 ± 55mm^2^, p<0.05) and SPHINX31 treated mice (516 ± 42.01/mm^2^, p<0.05) (Figure 3D). Similarly, the arteriolar density was significantly reduced in the ischemic gastrocnemius muscle of LysM-Wnt5a^GOF^ mice (4.32 ± 0.55/mm^2^) compared to WT mice (19.22 ± 2.03/mm^2^, p=0.004) and SPHINX31 treated mice (20.45 ± 4.22/mm^2^, p=0.002) (Figure 3E). Staining of gastrocnemius muscle from Wnt5a^GOF^, mice that had undergone ischemic ligation with an anti-VEGF-A_165_b antibody and IB4 showed that VEGF-A_165_b was seen in between the muscle fibres and around but not in the endothelial cells, but as a secreted protein its source was not clear. When co-staining with F4/80 (arrows), a macrophage marker, it was clear that macrophages in the Wnt5a^GOF^ mice expressed VEGF-A_165_b (inset). These results suggested that impaired revascularisation of LysM-Wnt5a^GOF^ may be dependent on VEGF-A_165_b expression controlled by SRPK1 activity.

**Figure 3.**
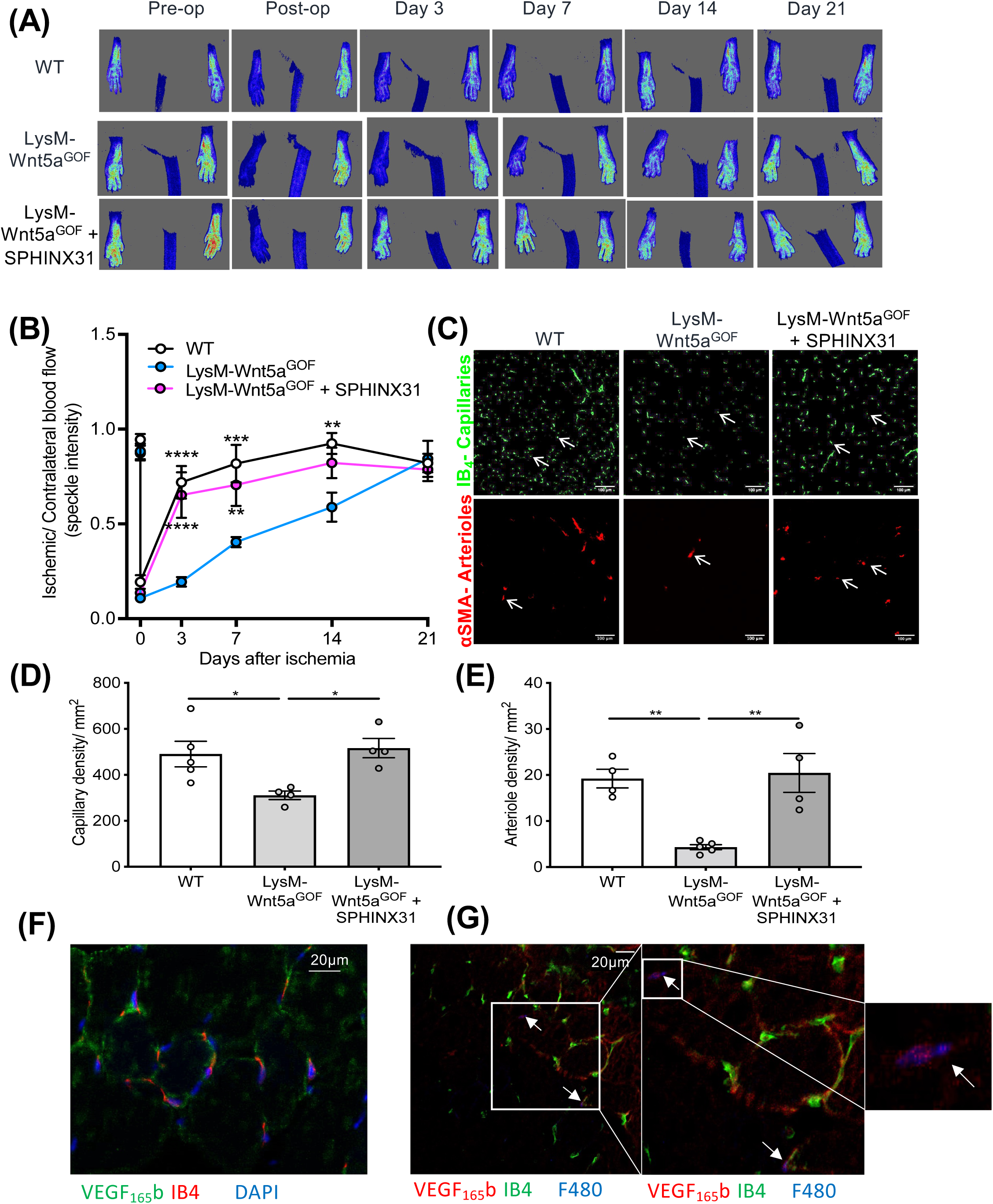
SRPK1 inhibition improves (and does not further impair) the blood flow recovery in LysM-Wnt5a^GOF^ mice. **(A)** LysM-Wnt5a^GOF^ mice underwent left femoral artery ligation followed by treatment with SPHINX31 biweekly (0.8mg/kg, i.p) or vehicle for the duration of the study. Blood flow to the paw was quantified using the laser speckle imaging system, as shown in the representative speckle images. **(B)** Quantitative analysis of the ischemic/ non-ischemic speckle intensity was calculated through to post-operative day 21. Data was analysed using a two-way ANOVA with post-hoc Bonferroni’s test (N=7-9/group). **(C)** Confocal images from the representative muscle sections of the gastrocnemius on post-operative day 21 stained with isolectin B4 for endothelial (green) and α-smooth muscle actin for vascular smooth muscle cells (red). Scale bar = 100 μm. **(D)** The capillary density was increased in LysM-Wnt5a^GOF^ treated with SPHINX31 compared with LysM-Wnt5a^GOF^ mice and matched the density in WT mice. Data was analysing using a one-way ANOVA with post hoc Bonferroni’s test (N=4-5/group). **(E)** The arteriole density was increased in LysM-Wnt5a^GOF^ treated with SPHINX31 compared with LysM-Wnt5a^GOF^ mice and matched the density in WT mice. (F) Wnt5a^GOF^ mice express VEGF-A_165_b in muscle. Triple staining for endothelial cells (IB4, red), VEGF-A_165_b (green) and nuclei (DAPI, blue). (G) Macrophages in Wnt5a^GOF^ mice express VEGF-A_165_b. Triple staining for endothelial cells (IB4, green), VEGF-A_165_b (red) and macrophages (F480, blue). Inset shows magnification of macrophage (blue cell) with VEGF-A_165_b expression (red). Data was analysing using a one-way ANOVA with post hoc Bonferroni’s test. (N=4-5/group). All results are shown as the mean ± SEM. *p<0.05, **p<0.01, ***p<0.001, ****p<0.0001.

### Monocytic-SRPK1 is responsible for impaired revascularisation in LysM-Wnt5a^GOF^ mice

Systemic treatment with an SRPK1 inhibitor could have many effects on many tissues. To determine whether monocytic-SRPK1 regulated the prolonged blood flow recovery in LysM-Wnt5a^GOF^ mice, we generated a double transgenic, monocyte specific LysM over-expressing, SRPK1 knockout mouse model (LysM-Wnt5a^GOF^:SRPK1^MoKO^, Figure 4A). These mice had no expression in CD11+ monocytes isolated from the spleen, or in bone marrow derived monocytes (Figure 4B), indicating a myeloid lineage deletion of SRPK1. These mice were subject to hind limb ligation, and quantitative analysis of the speckle images revealed that revascularisation in LysM-Wnt5a^GOF^ littermates had impaired flow compared to littermate control wild type mice on post-operative days 3 (0.19 ± 0.03 vs 0.48 ± 0.07, p=0.0034) and 7 (0.40 ± 0.03 vs 0.68 ± 0.07, p=0.004), which was rescued on day 7 in LysM-Wnt5a^GOF^:SRPK1^MoKO^ mice (0.65 ± 0.08, p=0.03) (Figure 4C-D). There was a moderate, but not statistically significant reduction in the capillary density in the ischemic gastrocnemius of LysM-Wnt5a^GOF^ mice compared with wild type which was partially but not significantly reversed by the SRPK1^MOKO^ (Figure 4E-F). However, the impaired arteriolar density in the ischemic gastrocnemius muscle of LysM-Wnt5a^GOF^ mice (4.32 ± 0.55/mm^2^) was significantly reversed in the LysM-Wnt5a^GOF^:SRPK1^MoKO^ mice (13.15 ± 2.48/mm^2^, p=0.007) and was comparable to WT mice (12.75 ± 1.60/mm^2^, p=0.009) (Figure 4G). To confirm knockout of SRPK1 we stained muscles for macrophages and SRPK1 (Figure 4H). Figure 4I shows that there was a significant reduction in SRPK1 positive F480 macrophages in the SRPK1^MoKO^ knockout mice compared with wild type, Wnt5a^GOF^ and Wnt5a^GOF^ mice treated with SPHINX31. Collectively, these results show that impaired revascularisation is dependent on monocytic-SRPK1 in LysM-Wnt5a^GOF^ mice.

**Figure 4.**
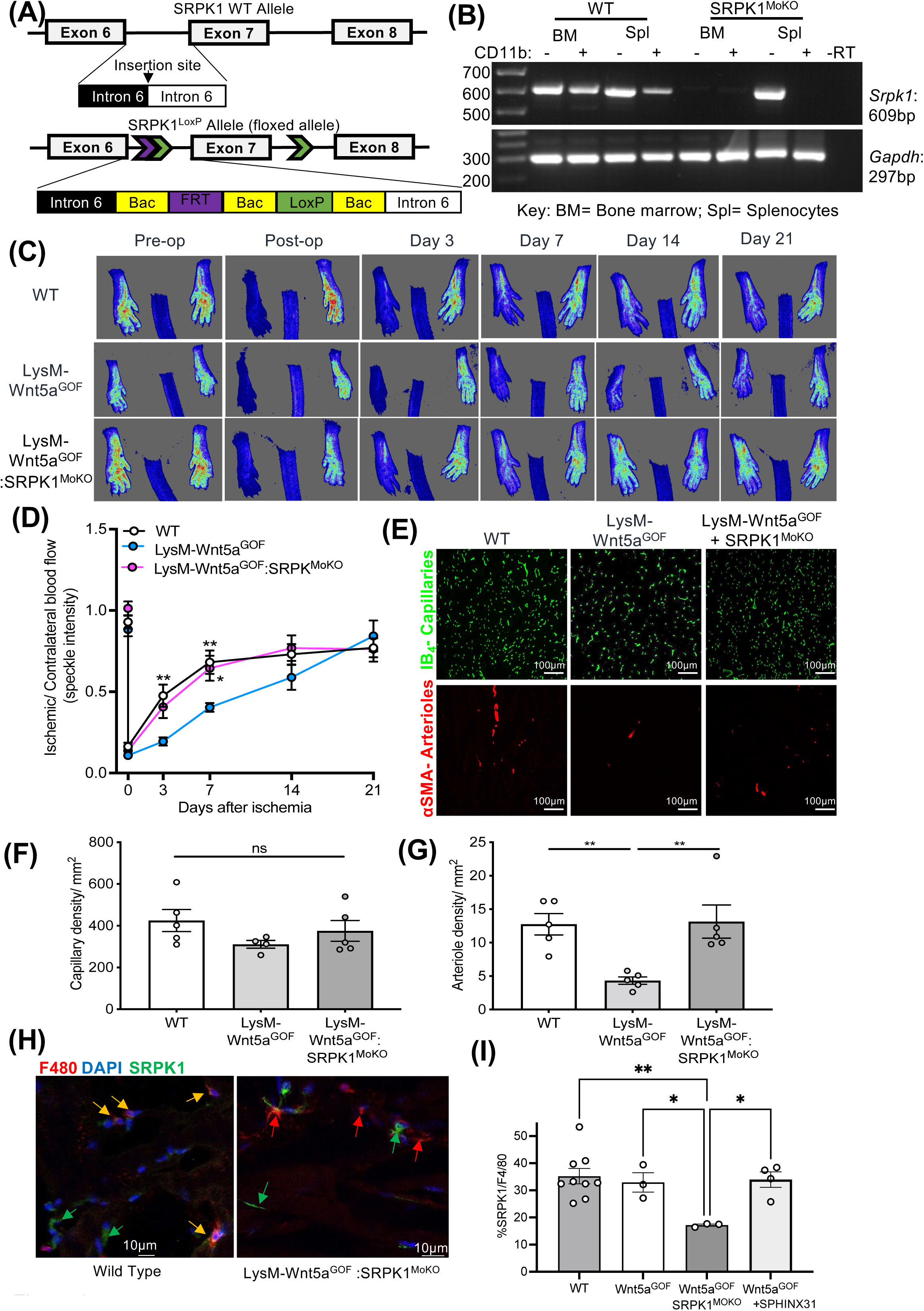
Impaired revascularisation in LysM-Wnt5a^GOF^ mice is controlled by monocyte specific SRPK1 activity *in vivo*. **(A).** Generation of the SRPK1 conditional knockout. Homologues recombination was used to insert LoxP sites either side of exon 7 of SRPK1. This results in deletion of exon 7 and loss of SRPK1 RNA presumably due to nonsense mediated decay. **(B)** CD11b^+^ Monocytes were extracted from mouse spleens and bone marrow from LysM-Cre: SRPK1^loxp/loxp^ mice (SRPK1^MoKO^) or wild type littermates (Cre negative mice) and subjected to PCR for SRPK1. In wild type mice SRPK1 was expressed in CD11b^+^ and CD11b^-^ cells from both bone marrow and spleen, but was only expressed in CD11^-^ cells from spleen in the SRPK1^MoKO^. **(C)** WT, LysM-Wnt5a^GOF^ and LysM-Wnt5a^GOF^: SRPK1^MoKO^ mice underwent left femoral artery ligation Blood flow to the paw was quantified using the laser speckle imaging system, as shown in the representative speckle images. **(D)** Quantitative analysis of the ischemic/ non-ischemic speckle intensity was calculated through to post-operative day 21. Data was analysed using a two-way ANOVA with post-hoc Bonferroni’s test (N=7-14/group). **(E)** Confocal images from the representative muscle sections of the gastrocnemius muscle on post-operative day 21 stained with isolectin B4 for endothelial (green) and a-smooth muscle actin for vascular smooth muscle cells (red). Scale bar = 100 μm. **(F)** The capillary density in WT, LysM-Wnt5a^GOF^ and LysM-Wnt5a^GOF^: SRPK1^MoKO^ mice was the same on post-operative day 21. Data was analysing using a one-way ANOVA with post hoc Bonferroni’s test (N=4-5/group). **(G)** The arteriole density was increased in LysM-Wnt5a^GOF^: SRPK1^MoKO^ mice compared with LysM-Wnt5a^GOF^ mice and matched the density in WT mice. **(H)**. Staining of gastrocnemius wild type or LysM-Wnt5a^GOF^:SRPK1^MoKO^ mice for SRPK1 (green) and F4/80 (red). Double stained cells (yellow arrows) were found predominantly in the wild type animals. **(I)**. Quantification of SRPK1-F4/80 double staining. Data was analysed using a one-way ANOVA with post hoc Bonferroni’s test (N=5/group). All results are shown as the mean ± SEM.*p<0.05, **p<0.01.

### SRPK1 knockout reverses impaired blood flow recovery in diet-induced obese mice

Previous studies have confirmed that mice fed on a HF/HS diet have an increase in Wnt5a expression in the ischemic gastrocnemius, which was co-localised to the macrophages(5). Therefore, we hypothesised that diet-induced obese mice would have impaired blood flow recovery, which would be reversed with SRPK1^MoKO^. Quantitative analysis of the representative speckle images in Figure 5A revealed that the level of the ischemia induced in the left hind limb of HF/HS-SRPK1^MoKO^ mice (0.12 ± 0.01) was more severe than that in HF/HS-WT mice (0.23 ± 0.03, p=0.02) (Figure 5B). Therefore, the flow was normalised to post-operative day 0, which revealed that rate of revascularisation was more rapid in obese-SRPK1^MoKO^ mice when compared to WT mice on post-operative days 14 (5.21 ± 0.41 vs 3.29 ± 0.41, p=0.002), 21 (5.30 ± 0.40 vs 3.55 ± 0.41, p=0.003) and 28 (6.29 ± 0.51 vs 3.73 ± 0.46, p<0.0001) (Figure 5C). Staining of capillaries and arterioles (Figure 5D) showed that HFHS-SRPK1^MoKO^ mice had a higher capillary density and arteriolar density compared to HFHS wild type mice after ischemia (Figure 5E).

**Figure 5.**
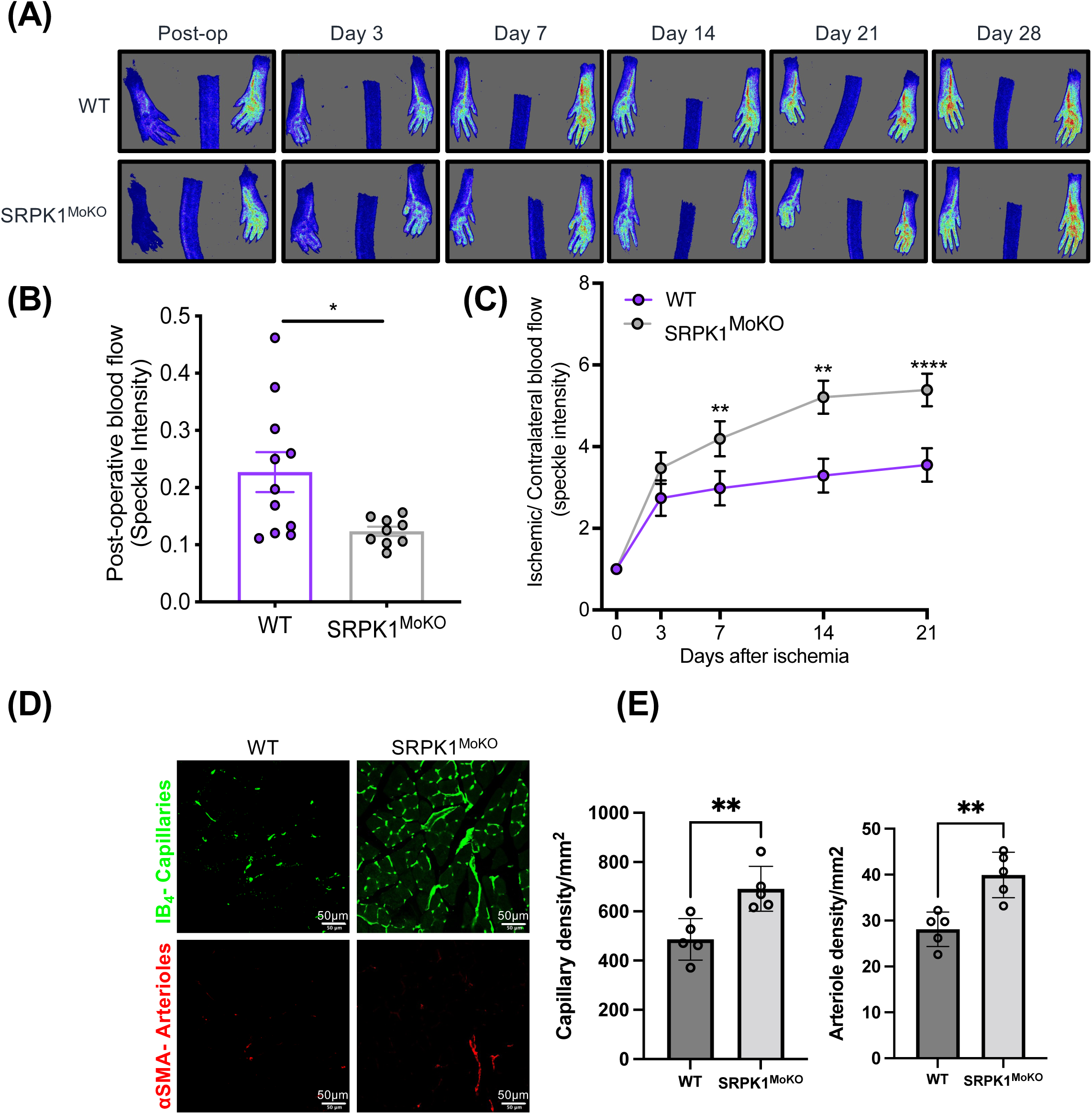
Blood flow recovery is reversed in SRPK1^MoKO^ mice fed on a HF/HS diet. **(A)** WT and SRPK1^MoKO^ mice were fed on a HF/HS diet underwent left femoral artery ligation Blood flow to the paw was quantified using the laser speckle imaging system, as shown in the representative speckle images. **(B)** Quantitative analysis of the ischemic/ non-ischemic speckle intensity indicates that the level of ischemia induced on post-operative day 0 was greater in SRPK1^MoKO^ mice compared to WT. Data was analysed using an unpaired t-test (N=9-11/group). **(C)** Data was further normalised to post-operative day 0 of each respective mouse in each group through to post-operative day 21. Data was analysed using a two-way ANOVA with post hoc Bonferroni test (N=9-11/group). **(D)** Confocal images from the representative muscle sections of the gastrocnemius muscle on post-operative day 21 stained with isolectin B4 for endothelial (green) and a-smooth muscle actin for vascular smooth muscle cells (red). Scale bar = 50 μm. **(E)** The capillary density in ischemic muscle of SRPK1^MoKO^ mice was significantly greater than wild type controls in obese animals. The arteriole density in obese mice was increased in ischemic muscle of SRPK1^MoKO^ mice compared with wild type controls. Data was analysing using an unpaired t-test (N=4-5/group). All results are shown as the mean ± SEM. *p<0.05, **p<0.01, ****p<0.0001.

To determine whether the phenotype of macrophages in the ischemic was changed by SRPK1 knockout or inhibition we co-stained tissues from the wild type, Wnt5a^GOF^, Wnt5a^GOF^-SRPK1^MoKO^ and Wnt5a^GOF^ treated with SPHINX31 (Figure 6) with the macrophage marker F480 and the subtype markers iNOS (M1 type macrophages, Figure 6A) or CD206 (M2 macrophages, Figure 6B). Staining showed that both types were present in ischemic tissues from wild type, Wnt5a^GOF^, Wnt5a^GOF^-SRPK1^MoKO^ and Wnt5a^GOF^-mice treated with SPHINX31. Counting of macrophages showed that there was a significant reduction in M1 macrophages in Wnt5a^GOF^-SRPK1^MoKO^ and Wnt5a^GOF^-SPHINX31 treated animals than in Wnt5a^GOF^ animals (Figure 6A,C) with a small but consistent increase in M2 macrophages (Figure 6B,D). This resulted in a significant reduction in the ratio of M1 to M2 macrophages in tissues where SRPK1 had been knocked out or inhibited (Figure 6E). Interestingly, F4/80 positive vessels were associated with blood vessels, but M2 macrophages appeared to be more associated with vessels resembling arterioles (Figure 6F) than venules or lymphatics. These results suggest that SRPK1 could be regulating the phenotype of macrophages leading to a more regenerative, pro-arteriogenic and less inflammatory phenotype.

**Figure 6.**
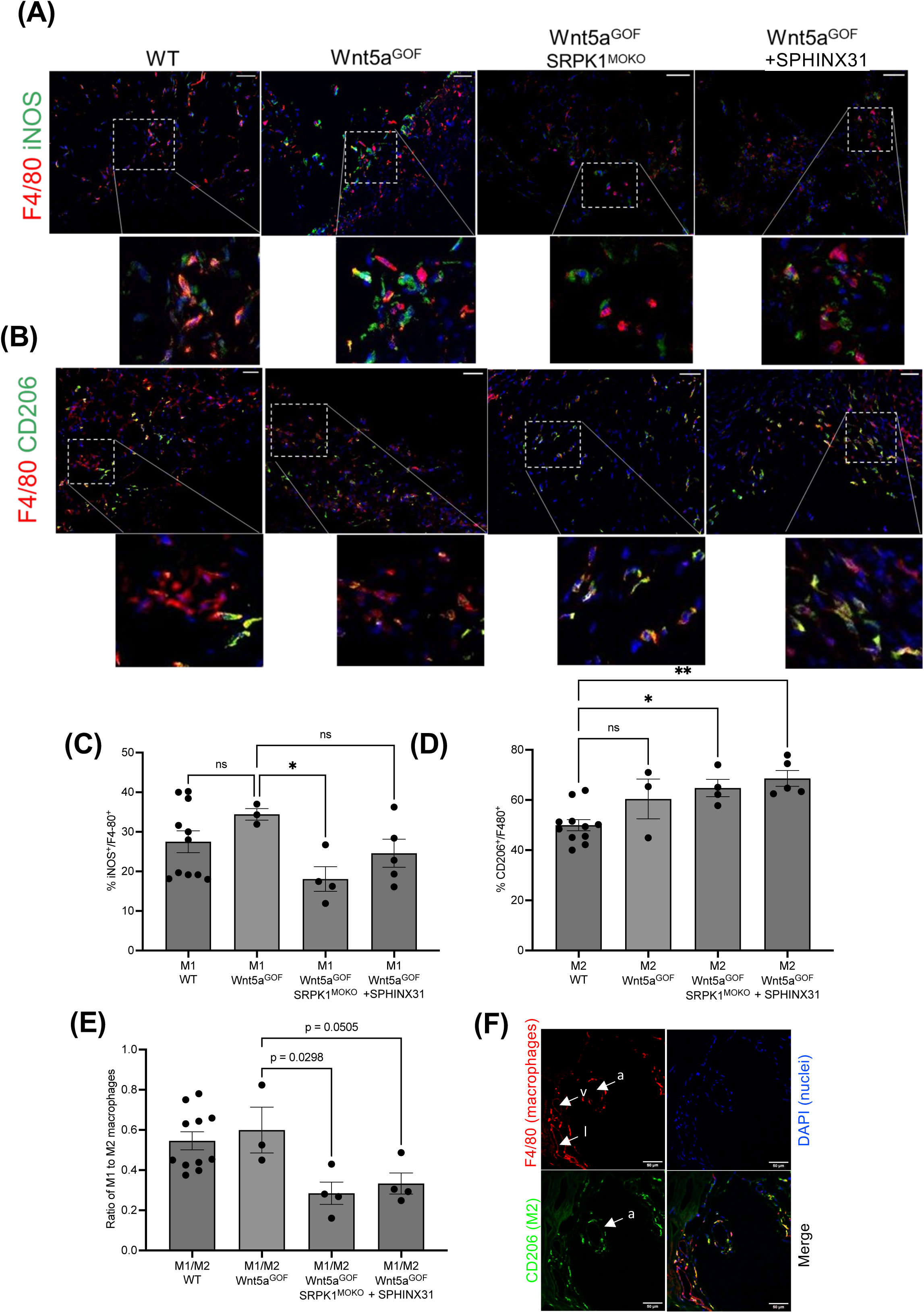
SRPK1 Knockout or inhibition alters macrophage phenotype. (A) Representative muscle images of M1 macrophages (red F4/80^+^ and green iNOS^+^) and (B) M2 macrophages (red F4/80^+^ and green CD206^+^) from wild type, Wnt5a^GOF^, WNt5a^GOF^-SRPK1^MoKO^ and Wnt5a^GOF^ treated with SPHINX31. Nuclei stained with 4’,6-diamidino-2-phenylindole (DAPI, blue). Each image presented with magnified field to highlight the phenotype of macrophage. Scale bar= 50 µm. (C) Quantification of macrophages expressing M1 phenotype (iNOS^+^) and (D) M2 phenotype (CD206^+^) respectively. Results are presented as percentage of double-positive macrophages from total macrophages (F4/80^+^). (E) Graph presenting M1/M2 ratio. Data was analysing using a one-way ANOVA with post hoc Bonferroni’s test. All results are shown as the mean ± SEM.*p<0.05, **p<0.01. (F) Staining of gastrocnemius muscle from ischemic wildtype mouse for CD206 (green) and F4/80 (red) showing accumulation of M2 macrophages around arterioles (a) but not venules (v) or lymphatics (l).

## Discussion

We show here that monocytes control VEGF splicing to be able to regulate angiogenesis through SRPK1 using both transgenic models driving VEGF splicing to the anti-angiogenic VEGF-A_165_b isoform, pharmacological and genetic manipulation of SRPK1 activity or expression, and human monocytes from patients with PAD. The implications of the finding are consistently that SRPK1 inhibition switches monocytes away from an anti-angiogenic phenotype, and in both the human monocyte *ex vivo* assay and the mouse *in vivo* assay, this is sufficient to induce behaviours associated with collateral formation (endothelial cell migration, new capillary and new arteriole formation).

We show in human cells that monocytes, when co-cultured with endothelial cells secrete the anti-angiogenic isoforms VEGF-A_165_b, as shown by the use of a neutralising antibody to VEGF-A_165_b. This antibody is raised against the C-terminal tail of VEGF-A_165_b and is effective in detecting VEGF-A_165_b and VEGF-A_189_b and the readthrough isoform VEGF-Ax(37). We have previously shown that this antibody has neutralising properties in the mouse models described here, which do not express VEGF-Ax(5). Treatment with an SRPK1 inhibitor increased VEGF-A_165_a from 0 to 40pg/ml, and reduced VEGF-A_165_b from 275 to 175pg/ml in the monocyte cell lysate. Interestingly, monocytes secrete sufficient VEGF-A_165_b to inhibit endothelial cell migration, and the neutralising antibody to VEGF-A_165_b completely reverses the monocyte mediated inhibition. Given that these experiments include 1nM VEGF-A_165_b, we can assume that the concentrations of VEGF-A_165_b in the media need to be close to that to inhibit endothelial cell migration. The proportion of VEGF that is VEGF-A_165_b in the monocytes has changed from 100% to 80% with SRPK1 inhibition, but this is sufficient to allow migration to occur. This suggest that the concentration of VEGF-A_165_b in the media must be close to the threshold for inhibition of VEGF-A_165_a, (potentially 1nM VEGF-A_165_b:1nM VEGF-A_165_a), and that switching the splicing therefore switches that to 0.8nM VEGF-A_165_b:1.2nM VEGF-A_165_a, i.e. from 50% to 40% VEGF-A_165_b. This again indicates that the switch in splicing does not have to be very great to allow angiogenesis to happen. Given that the angiogenic drive in the ischemic tissue is from the hypoxic muscle the balance between VEGF-A_165_b and VEGF-A_165_a in the tissues is likely to be critical for the efficacy of a splicing switch.

There have been surprisingly few studies investigating the effect of monocytes on endothelial cells in transwell studies (removing the interfering effects of contact between the cells). It has previously been shown that monocytes from healthy individuals stimulated to differentiate into macrophages can produce angiogenic effects by production of VEGF-A, depending on the differentiation state(38) and that this is inhibited by treatment with IL4 as this enhances production of the inhibitory isoform of VEGFR1, soluble flt1(39). This suggests that control of splicing of angiogenic processes in monocytes is a process that depends on the differentiation state of the monocytes and the environment in which they find themselves.

We also show that in transgenic animals with enhanced Wnt5a monocyte signalling(5), and in animals in which this signalling is induced metabolically(4), and in which VEGF-A_165_b is upregulated in monocytes(5), there is impaired collateral formation, which is reversed by SRPK1 inhibition. This finding is in direct contrast to the well described (including by us) action of SRPK1 inhibition in many other cell types, and was highly surprising, as we have previously shown that SRPK1 inhibition switches splicing from the VEGF-A_165_a isoform to VEGF-A_165_b in epithelial cells (retinal pigmented epithelial cells(33), glomerular visceral epithelial cells(27), in neurons (dorsal root ganglion neurons(40), and SHSY5Y neurons(41)), and in cancer cells (melanoma(42), prostate cancer (31) and cholangiocarcinoma cells(43)), and others have shown it regulates VEGF splicing in renal carcinoma(44), melanoma and Hela cells(45) and lung alveolar epithelial cells(46). Thus it appears that the control of VEGF splicing is not fixed, but dependent on either cell type or cell environment. While we cannot say for certain which of these is deterministic, it is unlikely that the diabetic environment controls whether SRPK1 determines splicing to VEGF-A_165_b or VEGF-A_165_a as SRPK1 inhibition switches splicing to VEGF-A_165_b in diabetic retinal epithelial cells(30), but the data here suggests that it switches it to VEGF-A_165_a in diabetic monocytes. SRPK1 acts via phosphorylation of SR rich sequences in SR proteins such as SRSF1(47). While we have shown that SRSF1 is the key target for SRPK1 in epithelial cells, we have not shown that for monocytes, and it should be noted that SRSF2 has been implicated in control of VEGF splicing in lung cancer cells(48) and in monocytes(5), suggesting that potentially SRPK1 may act in monocytes by phosphorylation of SRSF2 or other SRSF proteins. Furthermore, SRSF1 and SRPK1 are both significantly upregulated in ischemic Sfrp5^-/-^ mice (Appendix Figure 1), strongly suggesting that in monocytes, Wnt5a signalling can change splicing factor protein expression and hence alter the role of SR protein kinase. The finding that SRPK1 knockout or inhibition changes macrophage phenotype further suggests that the relationship between SRPK1 and VEGF splicing in this case is not direct, and also hints that macrophage phenotype maybe dependent on alternative splicing, perhaps in a wider context than just VEGF. Further work is required to determine the exact mechanism linking Wnt5a activation with SRPK1 activity or the link between SRP1 activity and macrophage phenotype and VEGF-A splicing.

The regulation of angiogenesis by SRPK1 may also be more complex than just regulating the VEGF-A_165_b to VEGF-A_165_b ratios as alternative splicing of VEGFR1 spice variants has also been demonstrated by SRPK1 through SRSF3 (49). This mechanism can now also be investigated using the SRPK1 knockout mice described here. In addition, we have studied here relatively young adult mice and in humans peripheral ischemia is predominantly a condition that affects older adults.

Finally, we describe here for the first time a viable SRPK1 inducible knockout mouse. Previously, Wang et al have described a conditional SRPK1 with a deletion of exon 3, but this mouse line was lost early after its development, and no further information has been available(50). We made numerous attempts to recreate this line using exon 3 and exon 1 LoxP flanking, but this was unsuccessful, presumably due to disruption of regulatory element within the introns. The insertion of the LoxP sites flanking exon 7 however, resulted in a robust knockout when crossed with Cre Driver lines, and the parental line has a normal phenotype. According to the International Mouse Strain Resource, there are 44 strains of mice that have been generated as ES cells that have an SRPK1 gene trap, one that is a CRISPR-Cas9 exon 3 knockout, and one that has a loxP exon 3 flanking regions. However, these mice have never been published or validated. The phenotype of genome wide SRPK1 knockout in adult animals is not yet known, but previous experiments have shown that, as here, systemic administration of SRPK1 inhibitors does not cause any imminent or obvious phenotype. Further work is required to determine whether SRPK1 inhibition could be a therapeutic approach in these types of condition, although given the finding that SRPK1 inhibition works differently in monocytes than for instance podocytes or retinal epithelial cells might make it difficult to predict adverse events.

While we have driven the knockout here in monocytes by crossing the SRPK1-LoxP mouse with a monocyte specific driver line, it is clearly of interest to generate lines where SRPK1 is knocked out in tissues in which VEGF expression is controlled in the opposite direction, e.g. in retinal epithelial cells(33), retinal neurons(30), and in renal epithelial cells(27), and determine whether control of isoform expression can affect diabetic nephropathy or retinopathy. SRPK1 knockout has been shown to be embryonically lethal before E14.5, as it appears to affect heart development(50) so this mouse is an ideal tool to investigate the role of SRPK1 in adult physiology. Further work will show how tissue specific and development stage specific SRPK1 knockout affects tissue function, and also the effects of global SRPK1 knockout in adult tissues. It was of interest that the monocyte specific knockout of SRPK1 (which would have occurred in the myeloid lineage from birth onwards(51)) had impaired flow after ligation indicating that residual collateral formation during development was impaired in these mice suggesting that myeloid cell derived SRPK1 is involved in the development of blood vessels.

In summary we show here that splicing control is cell type dependent, that monocytic SRPK1 controls VEGF splicing and that targeting SRPK1 can enhance collateral formation in models of peripheral vascular disease

## Acknowledgements.

This work was funded by the MRC (MR/K020366/1, MR/K013157/1), The British Heart Foundation (PG/21/10796, PG/13/47/30337, TG/18/3/33773) and the BBSRC Doctoral Training Program (BB/M008770/1).

---

**Appendix Figure 1.**
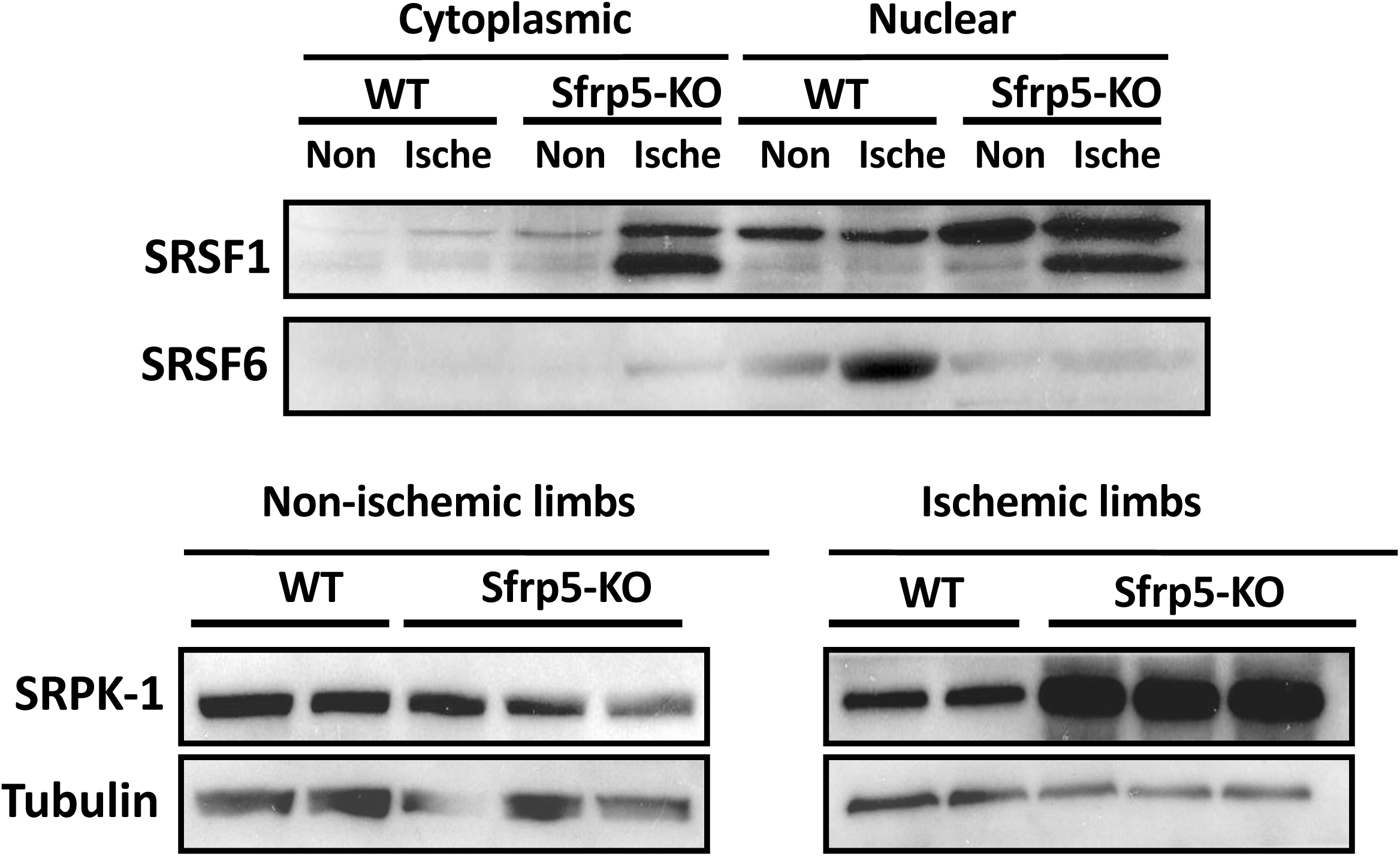
Muscle tissue was taken from Sfrp5^-/-^ or wild type littermate mice 7 days after ischemia was induced in the left hind limb. Cells were disaggregated and nuclear extract made and both nuclear and cytoplasmic extract subjected to immunoblotting for SRSF1 and SRSF6. Whole protein extract was subjected to immunoblotting for SRPK1.

